# A saturation mutagenesis approach to understanding PTEN lipid phosphatase activity and genotype-phenotypes relationships

**DOI:** 10.1101/255265

**Authors:** Taylor L. Mighell, Sara Evans-Dutson, Brian j. O’Roak

**Affiliations:** Neuroscience Graduate Program, Oregon Health & Science University, Portland, Oregon, USA.; Department of Molecular & Medical Genetics, Oregon Health & Science University, Portland, Oregon, USA.

## Abstract

Phosphatase and tensin homolog (PTEN) is a tumor suppressor frequently mutated in diverse cancers. Germline *PTEN* mutations are also associated with a range of clinical outcomes, including PTEN hamartoma tumor syndrome (PHTS) and autism spectrum disorder (ASD). To empower new insights into PTEN function and clinically relevant genotype-phenotype relationships, we systematically evaluated the effect of *PTEN* mutations on lipid phosphatase activity *in vivo*. Using a massively parallel approach that leverages an artificial humanized yeast model, we derived high-confidence estimates of functional impact for 7,244 single amino acid PTEN variants (86% of possible). These data uncovered novel insights into PTEN protein structure, biochemistry, and mutation tolerance. Variant functional scores can reliably discriminate likely pathogenic from benign alleles. Further, 32% of ClinVar unclassified missense variants are phosphatase deficient in our assay, supporting their reclassification. ASD associated mutations generally had less severe fitness scores relative to PHTS associated mutations (p = 7.16×10^-5^) and a higher fraction of hypomorphic mutations, arguing for continued genotype-phenotype studies in larger clinical datasets that can further leverage these rich functional data.

## MAIN TEXT

Recent large-scale exome sequencing studies have highlighted the abundance of protein-coding variation in the human population^1^. It remains challenging to predict variant pathogenicity and clinical outcomes, especially for genes with pleiotropic effects. With most rare variants private to a single family or individual, using traditional approaches to establish pathogenicity such as variant segregation within a pedigree or identification in independent patients is infeasible. Even for well-studied genes, hundreds of variants are currently defined as variants of uncertain significance (VUS). Moreover, purely computational approaches still suffer from high false positive rates^2^ and subjective interpretations that limit the clinical utility of these predictions.

To address these challenges for genes of clinical importance, one proposed approach is to prospectively measure the functional effects of all possible mutations, allowing these empirical data to be integrated into the clinical assessment of novel rare variants^3,4^. Historically, these types of functional assays have been conducted in a serial nature, which limits scalability, and often only within a portion of the protein of interest. While there are some notable examples of whole-gene brute force saturation mutagenesis, e.g., *TP53*^5^, new more scalable experimental paradigms are being developed that allow the functional dissection of the effects of thousands of genetic mutations in parallel^6^. These approaches leverage recent advances in DNA synthesis and sequencing technologies and have proven particularly valuable in understanding the effects of mutations in cancer-associated genes^7,8^.

With these issues in mind, we have developed a saturation mutagenesis approach to comprehensively assess the effect of nonsynonymous mutations on the lipid phosphatase activity of phosphatase and tensin homolog (PTEN). PTEN antagonizes the phosphoinositide 3-kinase (PI3K) signaling pathway through its lipid phosphatase activity toward the signaling lipid phosphatidylinositol (3,4,5)-trisphosphate (PIP_3_)^9,10^. In mice, loss of this activity increases tumor susceptibility in a dose dependent manner^11^. This observation led to a continuum model for PTEN’s role in cancer development, with the level of phenotypic severity tightly coupled to the level of lipid phosphatase activity^12^.

Germline *PTEN* mutations are associated with a diverse range of clinical outcomes, including autism spectrum disorder (ASD)^13–15^ and tumor predisposition phenotypes collectively known as PTEN hamartoma tumor syndrome (PHTS)^16–18^. Germline mutation carriers often share the common feature of increased head size or macrocephaly^19^. However, there is substantial variability in the neurological and tumor phenotypes present in these individuals. PHTS is an umbrella term that encompasses Cowden syndrome, Bannayan-Riley-Ruvalcaba-syndrome, and PTEN-related Proteus syndrome^20^. PHTS-affected individuals typically present with macrocephaly, hamartomatous polyps, and have an extremely high life-time risk of cancer^20^. *PTEN* mutations have been identified in macrocephaly cohorts of individuals with formal ASD diagnoses or developmental delay (DD)/intellectual disability (ID)^13,21,22^ as well as idiopathic ASD^14,23,24^.

It is currently impossible to predict the phenotypic outcome of a given *PTEN* mutation. Even predicting whether a *PTEN* mutation will have a pathogenic effect is still challenging. This is exemplified by the fact that a majority of missense variants (131/241, 54%) in ClinVar are considered VUS and the pathogenicity of 12 additional variants is inconsistently reported across laboratories. Recent evidence from functional assays on a limited number of mutations and using diverse models, including humanized yeast^25^, cultured human cells^26^, and *in vivo* mouse neurons^27^, suggest that mutations identified in individuals with ASD or DD without obvious PHTS features tend to have hypomorphic lipid phosphatase activity, while PHTS-associated mutations more frequently show complete loss of lipid phosphatase activity. Further supporting this hypomorphic hypothesis, the distributions of mutation types are consistent with ASD associated mutations being generally less severe, with reported missense mutations three to four times as common in ASD compared with PHTS^26,28^. These findings, as well as the established genotype-phenotype relationships for *PTEN* in cancer, led us to hypothesize that, at the population level, ASD-associated PTEN variants are hypomorphic compared to PHTS-associated PTEN variants.

To systematically test this hypothesis and improve our ability to interpret the functional effects of any *PTEN* mutation, we modified a previously validated humanized yeast model for massively parallel functional testing of the effects of *PTEN* mutations on lipid phosphatase activity *in vivo*^25,29^. Given that yeast do not signal through PIP_3_ dependent pathways^30^, this model system challenges PTEN protein variants to act on their preferred substrate in a cellular environment, but removes the confounding signaling and regulatory milieu present in mammalian cells. Accordingly, the model is more sensitive than *in vitro* assays in which PTEN dephosphorylates a water-soluble substrate^31^. The utility of the yeast model for measuring lipid phosphatase activity has been demonstrated through validation of mutation effects on downstream Akt1 activation in mammalian cells, exhibiting complete concordance for the variants tested^31^.

With this system, we analyzed the functional effect of 86% of all possible single amino acid alterations. Overlaying these data onto PTEN secondary and tertiary structures recapitulated many known or predicted structure-function and biochemical relationships but also revealed surprising patterns of mutational tolerance. We discovered that several residues within the catalytic pocket are surprisingly tolerant to mutation, and we identify residues that are critical for membrane interaction. Moreover, we demonstrate these functional fitness scores have clinical utility by showing that they can outperform *in silico-based* approaches in characterizing likely pathogenic and benign variants. Finally, we provide compelling support for the existence of germline *PTEN* genotype-phenotype relationships that should be further explored in larger longitudinal clinical cohorts.

## RESULTS

### Establishing a massively parallel functional assay for PTEN lipid phosphatase activity

We leveraged an artificial humanized yeast model in order to assess the relative phosphatase activity of PTEN variants^25,29^. In this system, the human PI3K catalytic subunit p110α (encoded by the *PIK3CA* gene) is expressed in *Saccharomyces cerevisiae* and artificially directed to the membrane by a C-terminal prenylation box^29^. At the membrane, p110α is able to catalyze the conversion of the essential pool of phosphatidylinositol (4,5)-bisphosphate (PIP_2_) to PIP_3_, which potently inhibits growth through cytoskeletal disruption^29^. Upon induction of gene expression, cells proliferate at a rate that is proportional to the ability of the PTEN variant to convert PIP_3_ to PIP_2_^31^. Co-expression of wild-type PTEN, but not catalytically dead mutants, e.g., C124S, catalyzes the reverse reaction, restoring the PIP_2_ pool and allowing the yeast to grow and survive (Fig. 1a). Moreover, growth rate provides a quantitative surrogate of lipid phosphatase activity with partial loss of function mutations showing intermediate growth phenotypes^25^.

**Fig. 1.**
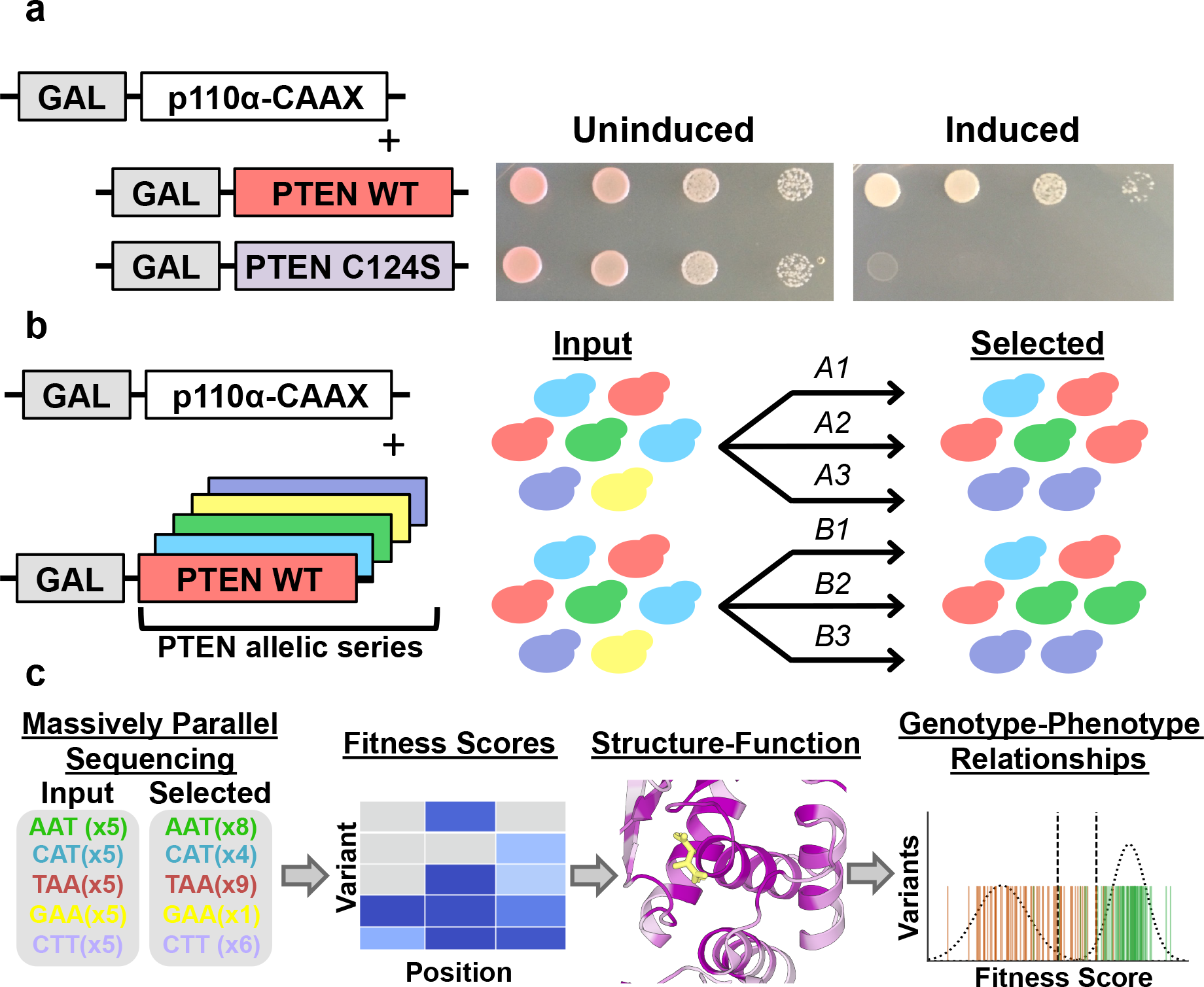
A framework for massively parallel functional testing of PTEN mutations. **a,** Humanized yeast model for evaluating the effect of PTEN mutations on lipid phosphatase activity. Exogenous expression of the catalytic subunit of human PI3K with a membrane-targeting prenylation motif (p110μ-CAA☓) in yeast is toxic. However, co-expression of human PTEN wild-type, but not catalytically-dead PTEN C124S, can rescue growth. Both genes are under the control of a galactose inducible promoter (GAL). **b-c,** Modifications to allow massively parallel variant assessment. **b,** We generated a comprehensive PTEN allelic series, introduced these variants into yeast en masse, and subjected them to p110μ-CAA☓-mediated selection in liquid culture. We performed two biological replicates, each consisting of three technical replicates. **c,** We collected input and selected timepoints and subjected these to deep sequencing. We used read counts to calculate fitness scores and used these scores to highlight structure-function insights as well as genotype-phenotype relationships.

We made several modifications to this system that allowed for massively parallel testing of preprogrammed mutations. First, to allow for parallel testing, rather than serial plating of single mutations, we modified the assay to support complex populations of PTEN-bearing yeast in liquid culture and sequencing as a readout of growth (Fig. 1b,c, Supplementary Fig. 1). We then introduced the yeast-preferred codon for each non-wild-type amino acid, stop codon, and single residue deletion at all *PTEN* codons en masse, utilizing a homologous recombination-based mutagenesis approach^32,33^ (Methods, Fig. 1b, Supplementary Fig. 2a, Supplementary Table 1). To allow direct sequencing of each mutagenized region, mutational space was separated into ~300 base-pair quadrants (Supplementary Fig. 2a).

We transformed two independent yeast populations with our mutagenesis library. Sequencing of naïve yeast libraries indicated that 95% of all intended mutations were present (Fig. 1b, Supplementary Fig. 2a). No position had less than 33% mutational coverage. Mutation dropout was largely confined to a single oligo pool in the C2 domain of the protein, which repeatedly performed poorly. We then performed selection experiments on these two independent yeast populations, each with three selection replicates (Fig. 1b). We calculated natural log-scaled and wild-type normalized fitness scores for each variant, along with standard error-based confidence intervals^34^ (Methods, Supplementary Fig. 2b). Score estimates were generated for 8,019 (95% of intended) *PTEN* nonsynonymous mutations and between mutational libraries fitness scores were highly correlated (Pearson’s r = 0.76, Table 1, Supplementary Fig. 3a,b, Supplementary Table 2). The distribution of fitness effects illustrates two major populations corresponding to likely damaging and wild-type-like mutations (Supplementary Fig. 3a). Based on low standard error or replicate concordance, scores for 7,244 mutations (86% of intended) were classified as high-confidence (Methods, Table 1, Supplementary Fig. 3c, Supplementary Table 2). Mutations were classified as wild-type like if their average fitness score was within the 95^th^ percentile (two-tailed) of observed synonymous mutations (Supplementary Fig. 3d). We identified 2,273 likely damaging mutations (31%), 4,872 wild-type-like mutations (67%), and 99 mutations that performed better than wild-type (1%) (Table 1). Among the likely damaging missense mutations, 1,243 (17%) fell within the observed distribution for programmed premature truncations (excluding C terminal tail), with the remainder having intermediate phenotypes in this assay.

**Table 1.**
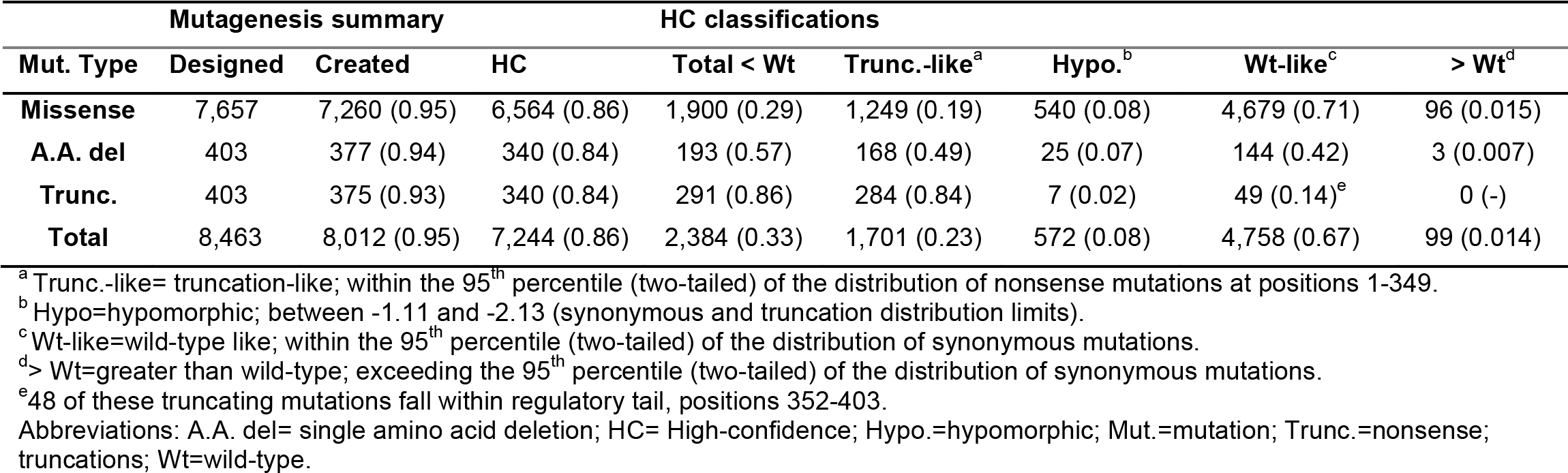
Summary of *PTEN* mutagenesis and high-confidence effect classifications.

### High-resolution mutation data reveal structure-function insights

Using the high-confidence data, we first analyzed structure-function relationships, including known or predicted functional domains. Our complete sequence function map recapitulates many known features of PTEN biochemistry. For example, early truncating mutations are uniformly damaging through the phosphatase and C2 domain, but are tolerated in the regulatory tail^31^ (Fig. 2a). Overlaying the median fitness score of each position onto the partial crystal structure of PTEN (including residues 7-285 and 310-353) reveals strong intolerance of positions in the phosphatase domain, especially those positions near the catalytic pocket (Fig. 2b). The median fitness scores are also correlated with evolutionary conservation (Spearman, ρ = 0.58, Supplementary Fig. 3e). When compared to positions in alpha helices and beta strands, unstructured positions are very tolerant to mutation (Supplementary Fig. 3f).

**Fig. 2.**
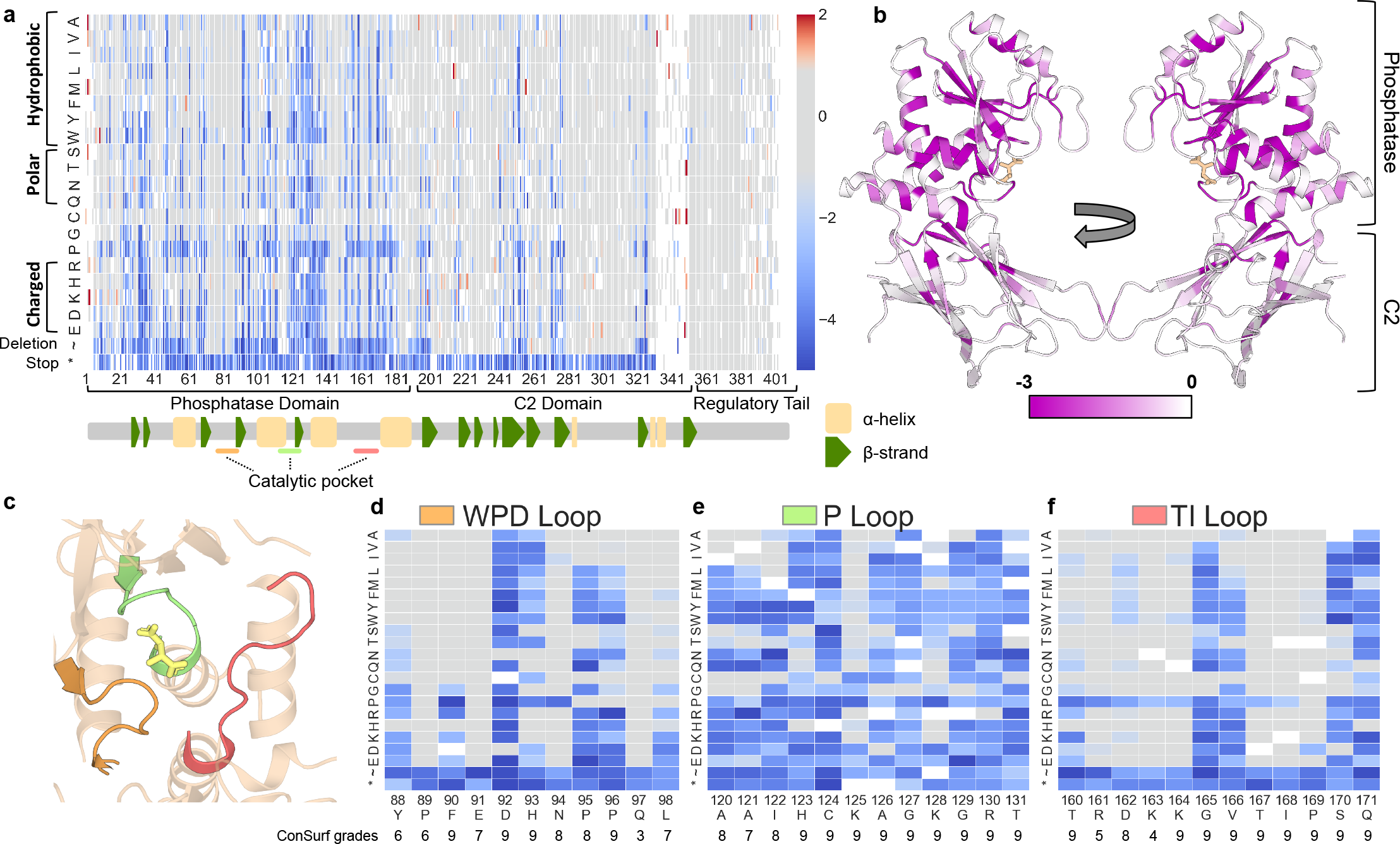
High-resolution map of the functional effects of PTEN mutations. **a,** Heatmap schematic showing High-confidence fitness scores for 7,244 PTEN missense, nonsense, or in-frame deletion mutations (86% of possible). Columns are each protein position and amino acids are listed in rows ordered according to biophysical characteristics. Variants with fitness scores within the 95^th^ percentile (two-sided) of synonymous wild-type like mutations are colored gray. Variants with fitness scores lower than the synonymous distribution are colored blue while variants with higher fitness scores are colored red. The major protein domains, as well as the secondary structure features are indicated in the track below the heatmap (a-helices as yellow rectangles and β-strands as green pentagons). **b,** Ribbon diagram of PTEN crystal structure with residues colored by average fitness score. Darker purple corresponds to more damaging scores, **c,** Ribbon diagram highlighting the crystal structure of the PTEN catalytic pocket, composed of the WPD (orange), P (green), and Tl-loops (salmon), **d-f,** The fitness scores of mutations at the residues composing the three catalytic pocket loops. Beneath each position is the Consurf grade (Methods), which represents the relative evolutionary conservation, with nine being the most conserved and one being the least conserved.

The catalytic pocket of PTEN is composed of the WPD, P, and TI loops (Fig. 2c). This motif has sequence homology to dual specificity protein phosphatases, especially within the signature motif (123-HC☓☓G☓☓R-130)^35^. R130 is a hot-spot for somatic cancer associated mutations with multiple different missense and truncations frequently reported^36^. We observed this critical position was intolerant to all mutations (Fig. 2e). Compared to other phosphatases, PTEN also has unique sequence features in order to accommodate the highly acidic and bulky PIP_3_ substrate. Residues H93, K125, and K128 impart a basic character on the pocket^35^, the importance of which is demonstrated by the mutational intolerance at these positions (Fig 2d,e). D92 is a critical residue for PTEN catalysis, but its exact role remains uncertain^25,37^. We find that the only substitution with wild-type like activity is asparagine. Additionally, the PTEN catalytic pocket is larger compared to other dual specificity phosphatases^35^. The Cowden-associated G129E mutation has been shown to abolish lipid phosphatase while preserving protein phosphatase activity^38^. Our data show G129 is intolerant to all mutations except to alanine and serine, the two next smallest amino acids (Fig. 2e). Unexpectedly, despite their presence in the catalytic pocket, several residues in the WPD and TI loops are highly tolerant to mutations (Fig. 2d,f), highlighting the power of functional data to delineate truly functional from non-functional alterations within highly conserved protein domains.

### PTEN associates with the plasma membrane through multiple domains

A PIP_2_ binding motif in the phosphatase domain (residues 6-15) is rich in positively charged amino acids and allosterically promotes catalysis upon PIP_2_ binding^39,40^. An additional positively charged residue, R47, contributes to this interaction^41^. Our data suggest that R15, K13, and R47 are the most critical of the positively charged residues in this motif^42^ (Supplementary Fig. 4a). Additionally, an intramolecular regulatory interaction between the C-terminal tail and the phosphatase domain is controlled by phosphorylation at four sites in the tail, in mammalian cells^43^. We find that individual phosphomimetic substitutions at these sites are insufficient to decrease activity in our assay (Supplementary Fig. 4b).

Protein positions cluster into stereotyped patterns of mutational sensitivity. In order to identify patterns of mutational sensitivity among PTEN positions and amino acid substitutions, we performed hierarchical clustering with all positions at which we measured effects of all missense substitutions (including high and low confidence, n = 326, Fig. 3a). We found that positions clustered into two major clades, corresponding to positions broadly tolerant/intolerant to proline or highly sensitive positions. We identified solvent exposure as a highly discriminatory feature between sensitive and tolerant clades, with 80/88 (91%) positions in the sensitive clade being in buried positions, while only 44/170 (26%) are buried in the tolerant clade (Fig. 3a). The tolerant clade splits into two major groups with a sub-clade broadly tolerant to all substitutions (beige) and a second sub-clade where positions are either sensitive to proline alone or proline and hydrophobic residues (purple). The proline sensitive positions generally are part of secondary structures that are not buried in the hydrophobic core (Fig. 3a). The sensitive clade positions split into three groups (green shaded sub-clades), which differ in their tolerance to charged, polar, or hydrophobic residues. The dark green clade represents the most constrained positions, and includes position 92, 123, 124, and 130; all of which are in the catalytic pocket and critical for catalysis. Overlaying the sub-clade assignment of each position onto the crystal structure highlights the intolerance of mutations within the hydrophobic core of the phosphatase domain. Many of the solvent exposed positions in the C2 domain are tolerant to mutation (Fig. 3b).

**Fig. 3.**
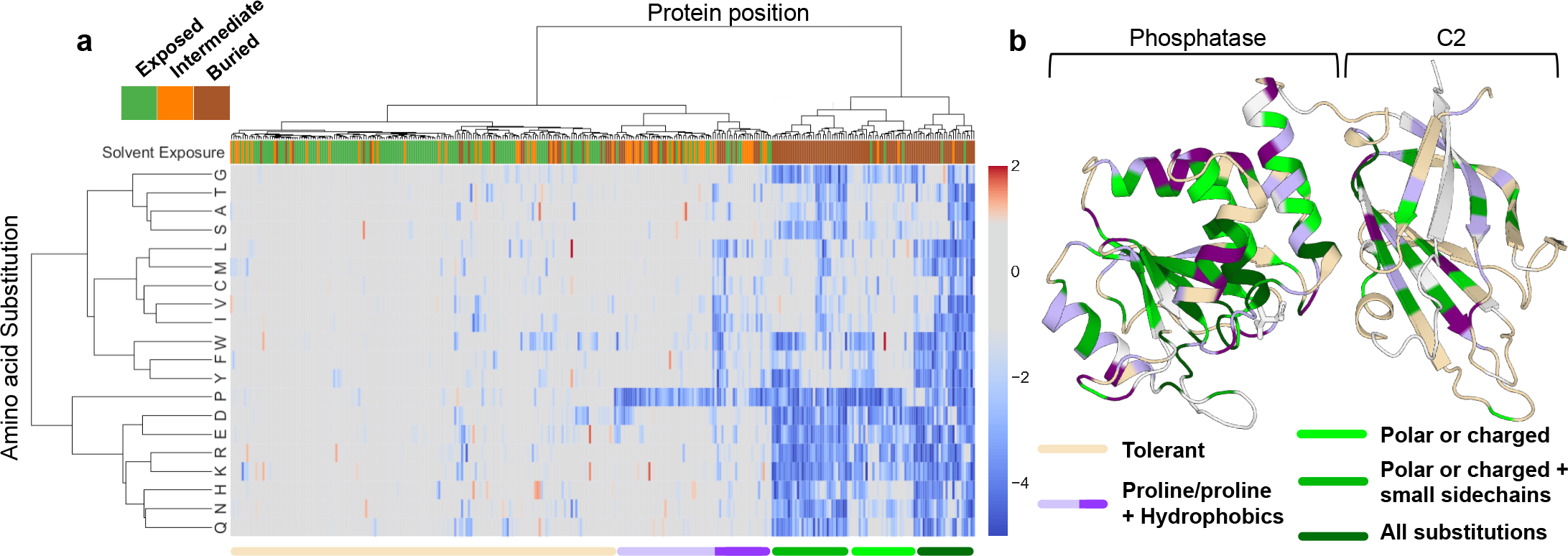
Hierarchical clustering reveals patterns of mutational tolerance among protein positions and amino acid substitutions. **a,** Hierarchical clustering of the 326 sites with all missense mutations measured. Clustering was performed by positions and amino acid substitutions (positions are columns and amino acid positions are rows). Overlaid on this heatmap is a top track showing the solvent exposure of each position in the crystal structure (1D5R), with solvent exposed positions colored green, intermediate positions orange, and buried positions brown. We identified two major clades, which partitioned into five sub-clades with prevailing characteristics indicated and represented in the bottom track. We further divided the purple clade to reflect major differences in mutational tolerance. **b,** Ribbon diagram of PTEN crystal structure with residues colored according to clade assignment.

Clustering by amino acid substitutions recapitulated known functional relationships with proline correlated poorly with other substitutions (Fig. 3a). We sought to leverage these patterns of correlation to predict the fitness scores of mutations that were not present in our mutagenesis library or that were lowconfidence^44^. We developed a heuristic for using only the most closely correlated observed substitutions^45^ at the site of interest to compute an “informed position average” (Supplementary Fig. 5a). We combined this with several other prediction based, evolutionary, and biophysical features to train and test a random forest regression algorithm on our high-confidence measurements^44^ (Methods, Supplementary Fig. 5b,c, Supplementary Table 6). We used 10-fold cross validation to confirm that this approach can predict unseen data with high confidence (Pearson’s r = 0.80, Supplementary Fig. 5e). We further performed a downsampling analysis to assess the expected accuracy of imputing scores at different levels of saturation, finding that reductions of 10-20% (65.8-74% of saturation) achieve similar performance (Supplementary Fig. 5f). Finally, we generated imputations for all variants that were absent from our library or measured with low-confidence (Supplementary Fig. 6).

### Fitness scores discriminate between likely pathogenic and benign alleles

To determine if our empirically determined fitness scores were informative for discriminating between germline likely pathogenic and benign alleles, we collected germline missense mutations reported as pathogenic or likely pathogenic from ClinVar^46^ and rare variants from gnomAD^1^ (excluding R173H and K289E, which are reported pathogenic in ClinVar, Methods, Supplementary Tables 3,4). Fitness scores alone discriminated pathogenic from benign germline alleles (Fig. 4a). We found that the F_0.5_ score, which weights predictive value (PPV) over sensitivity, reaches its maximum at a cutoff based on the synonymous distribution (<= ‐1, ~95^th^ percentile, PPV = 0.93, sensitivity 0.83), and outperforms several *in silico* mutation effect prediction algorithms (Fig. 4c). PPV was maximized (0.98) at a more conservative cutoff based on the 95^th^ percentile of the truncation distribution, but with reduced sensitivity (0.60) (Fig. 4a,c). Given the high PPV of our scores, we evaluated distribution of fitness scores among ClinVar missense VUS (Fig. 4b). We found that 21/127 (17%) VUS with high-confidence data met the strict truncation-based cutoff and 41/127 (32%) met the synonymous cutoff, suggesting that fitness scores could be used to reclassify a major fraction of VUS.

**Fig. 4.**
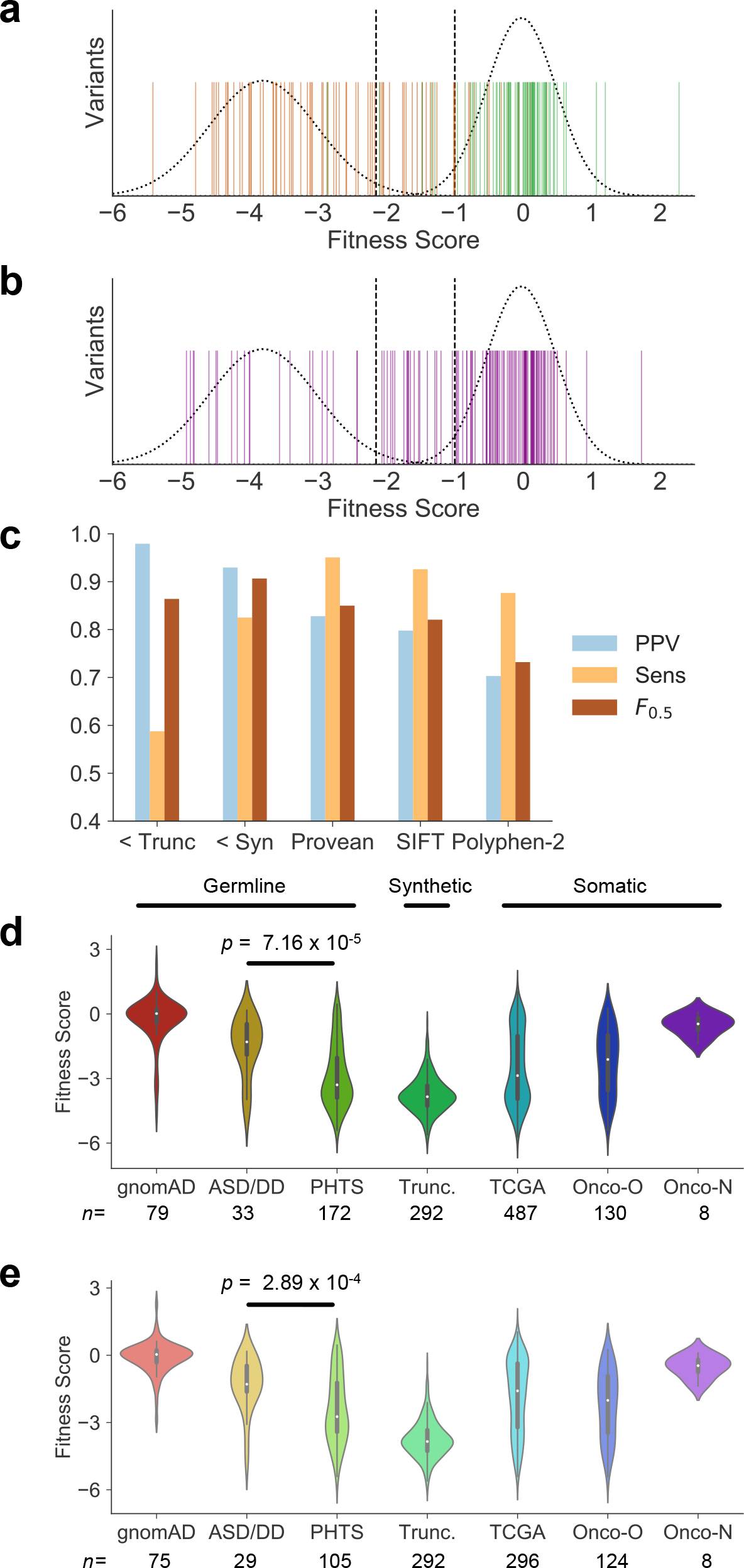
Fitness scores discriminate between likely pathogenic and benign variants and support genotype-phenotype relationships. **a,** Fitness scores for missense variants considered pathogenic or likely pathogenic in ClinVar (orange) and putatively benign variants from gnomAD (green). Dashed lines at ‐2.15 and ‐1 represent the approximate 95% two-tailed distribution of truncations (before regulatory tail) and synonymous mutations, respectively. **b,** Fitness scores of ClinVar VUS (purple), with truncation and synonymous distributions and 95% limits. **c,** To test the ability of fitness scores to discriminate likely pathogenic from likely benign missense mutations, we calculated positive predictive value (PPV), sensitivity, and F_0_.s scores for our fitness scores (“< Trunc” represents the threshold at ‐2.15, “< Syn” represents the threshold at ‐1). In these tests, a true positive represented a ClinVar pathogenic allele having a fitness score less than or equal to the threshold. We compared the performance of the fitness scores at these two thresholds with *in siiico* pathogenicity predictors for missense mutations (Methods). **d,** Fitness scores of all curated mutations associated with the indicated phenotype (Methods). ASD/DD= autism spectrum disorder/developmental delay, PHTS= PTEN hamartoma tumor syndrome, TCGA= The Cancer Genome Atlas, Onco-0= OncoKB mutations considered “oncogenic” or “likely oncogenic”, Onco-N= OncoKB mutations considered “likely neutral.”

*PTEN* mutations are extremely frequent in somatic cancer. We extracted nonsynonymous mutations from The Cancer Genome Atlas (TCGA) and observed a multimodal and wide distribution of fitness scores (Fig. 4d,e, Supplementary Table 5). This is likely due to the presence of both driver and passenger mutations in these data. Similar to the germline analysis, to test if fitness scores could discriminate somatic mutations that are likely pathogenic, we evaluated mutations from Onco-KB, a precision oncology database with expert annotation of somatic mutations^47^ (Supplementary Table 5). We found that fitness scores of *PTEN* mutations considered “oncogenic” or “likely oncogenic” were substantially more negative than those considered “likely neutral.” Of the missense likely oncogenic, 86/124 (69%) and 56/124 (45%) were below the synonymous and truncation thresholds, respectively. In contrast, of the 8 variants considered likely neutral (all missense), only one (A121V) had a fitness score marginally below the synonymous cutoff (fitness score, ‐1.3). Taken together, these findings emphasize the ability of empirically determined fitness scores to discriminate between pathogenic and benign human alleles, in both the germline and somatic setting.

Finally, we evaluated potential genotype-phenotype relationships for germline *PTEN* mutations. We first compared the fitness scores of *PTEN* mutations associated with various clinical presentations acquired from multiple sources (Methods, Fig. 4c, Supplementary Table 5). We found that, as a population, fitness scores of nonsynonymous mutations exclusively reported in ASD/DD cohorts were less severe than PHTS-associated mutations (Mann-Whitney U-test, two-sided, p = 7.16 × 10^-5^). Comparing only the missense we found that this significant difference persists (Mann-Whitney U-test, two-sided, p = 2.89 × 10^-4^), indicating that the mutation type alone does not drive these differences. We found 12/29 (41%) and 21/105 (20%) of the ASD and PHTS missense mutation fell within the hypomorphic activity range, respectively. Overall, these data provide strong support for the hypothesis that ASD/DD associated mutations often retain hypomorphic PTEN phosphatase activity.

## DISCUSSION

Massively multiplexed functional assays represent a promising approach to understanding the effect of mutations on protein function, which can provide immediate insights into structure-function relationships and clinical interpretation. Modifying a humanized yeast assay that uses growth to readout relative phosphatase activity, we were able to assess the functional effects of human *PTEN* mutations on a massive scale. Our approach yielded high-confidence measurements of 86% of the possible single residue nonsynonymous mutations. A limited number of human proteins have been subjected to full length massively multiplexed functional assessment, and very few have been assayed at the depth we achieved^7,8,44,48–52^. Similar approaches could be used with this model to the study of various aspects of the PI3K/Akt pathway at scale, including mutations in PIK3CA/B (p110μ/β)^31^, PIK3R1 (p85a)^53^, and AKT1^54^ as well as drug screening for PIK3CA inhibitors^55^.

Several features of the data support the validity of these function estimates and their relevance to human health. We observed high correlation between biological replicates and recapitulated known features of PTEN function. For example, the set of early terminating mutations confirm that the minimal catalytic unit includes the phosphatase and C2 domains, but not the C-terminal tail^31^. Likewise, we found that position C124, which takes part directly in phosphatase catalysis, and position R130, which is a hotspot for cancer mutations, are completely mutation intolerant. Additionally, we found that mutations are not well tolerated within the loops forming catalytic pocket or residues mediating interactions with PIP_2_. Finally, we found that proline was the most damaging substitution, consistent with a recent meta-analysis of massively multiplexed experiments^45^ and decades of biochemistry^56^.

While the humanized yeast system faithfully reports on intrinsic lipid phosphatase activity, mutations that functionally disrupt protein-protein interactions, subcellular localization, post-translational modifications, or function through a dominant negative mechanism^57^ in mammalian cells will not be captured. PTEN has relatively low thermostability^58^, and protein destabilization is a known mechanism for PTEN loss-of-function^26,59^. A concurrent functional screen assaying protein stability found ~1/4^th^ of mutations alter steady state stability^51^. Six mutations that destabilized PTEN in breast cancer cell lines also decreased steady state abundance in this yeast model^31^, suggesting that mutations affecting thermostability will be detected in our screen. However, our sensitivity to detect destabilizing mutations is unknown, as is whether mutations specifically altering the rate of proteasome-mediated degradation^60^ will be reported on. We believe that independently assaying these important factors at similar scale would provide useful complementary insights into PTEN function.

We discovered that approximately half of all positions in PTEN were broadly tolerant to substitutions, suggesting that they are not required for lipid phosphatase activity. While there is a degree of correlation between the median fitness score and the evolutionary conservation of each position, we identified positions within the highly conserved catalytic pocket and elsewhere in the protein that are highly tolerant to specific mutations. This is in apparent contradiction with PTEN’s high evolutionary conservation (99.75% identity between human and mouse^28^) and constraint in humans^1^. This suggests that many PTEN positions are potentially under selection due to phosphatase independent functions.

Our high-resolution mutation data empowered unique insights into PTEN biochemistry and structure. G129E is a well-known Cowden-associated mutation that disrupts lipid phosphatase activity while maintaining protein phosphatase activity^38^. We found that substitutions to alanine and valine are tolerated at this position, while mutations to bulkier residues are damaging. This suggests that there is a size limit for the amino acid that occupies this position. D92 matches the position of aspartic acid in the WPD loop of PTP1B, which acts as a general acid in the catalytic mechanism^37^. D92 is a critical residue in the PTEN catalytic pocket, but its role in the reaction mechanism remains uncertain^25,37,61^. Our data support previous findings that all mutations except D92N are damaging^25^. However, a D92N mutation has been reported in an individual with ASD indicating that it still may have a clinical effect^62^. Asparagine deamidation is a spontaneous, intramolecular reaction that can result in the conversion of asparagine to aspartic acid^63^. We propose that D92N could be showing wild-type like activity as a result of this reaction in our assay. In mammalian cells, this spontaneous conversion may not be sufficient to fully rescue PTEN activity.

Similar to previous studies^5,64^, we used hierarchical clustering to look for patterns amongst the positions and amino acid substitutions. We found that PTEN positions fall into a few stereotyped patterns of mutational tolerance and that a critical determinant of mutational tolerance is the relative solvent exposure of the position. These findings are consistent with a recent meta-analysis of similar experiments^65^. We leveraged the correlation amongst amino acid substitutions, along with several other features, to generate a random forest regression model that could accurately predict the fitness scores of unseen mutations and create a comprehensive functional map encompassing the effects of all possible single nonsynonymous mutations. To guide future studies of similar proteins, we performed a downsampling analysis of the training data and found that for similar accuracy, ~70% mutation saturation would likely be sufficient. Moreover, proline substitutions predict poorly and should be directly assayed.

A critical hurdle for the application of massively multiplexed functional assays is bridging the gap between molecular phenotype and human phenotype^66^. We found that fitness scores are able to discriminate between likely pathogenic and benign human alleles in both the germline and somatic condition. On this basis, we expect that these scores will be of tremendous clinical value for reclassifying VUS^4^ and also predicting the effects of private alleles that remain to be identified. A major question related to *PTEN* genetics is whether genotype-phenotype relationships can explain the heterogeneity in clinical presentation for carriers of germline mutations. Our comprehensive dataset provides strong evidence that the mutations associated with ASD/DD are hypomorphic for lipid phosphatase activity and are significantly more active than the mutations that lead to PHTS. This suggests that distinct biological mechanisms underlie the differential presentations, and understanding these differences will be critical to the eventual treatment of these disorders. While it is possible that these different mechanisms are the direct result of lipid phosphatase activity at the plasma membrane, ASD-associated mutations may specifically disrupt another of PTEN’s cellular functions^67,68^. Supporting this idea, some ASD-associated mutations are excluded from the nucleus and lead to neuronal hypertrophy, but this phenotype can be rescued by artificial direction to the nucleus^69^.

While massively parallel functional data is a significant advance for understanding function-specific mutation effects, further untangling complex genotype-phenotype relationships will require similar advances in clinical genetics databases with standardized descriptors of clinical presentations and symptoms^28^. Our study was limited by both the number of publicly available mutations and associated clinical information. Since there are no variants considered benign in ClinVar, we used PTEN variants in the gnomAD database as a proxy for likely benign mutations. While these mutations are on average wild-type like, we recognize that this is an imperfect approach and it is possible that some of the variants in gnomAD are pathogenic. We excluded variants that were only in ClinVar from our genotype-phenotype analysis because of their ambiguous annotation and lack of clinical data. For example, 17% of the pathogenic/likely pathogenic mutation submissions had no indicating condition provided and 36% of all missense entries use the ambiguous term “hereditary cancer-predisposing syndrome.” Requiring submitters to provide more information in a consistent way will maximize the utility of massively multiplexed functional data. Finally, it is still unclear if individuals ascertained for neurological phenotypes as children will have a higher risk to develop PHTS like or cancer presentations later in life^70^. Moving forward, large-scale sequencing efforts that permit longitudinal assessment as well as patient re-contact will be instrumental. A new initiative, SPARK, aims to partner with 50,000 individuals with ASD and their families to create the largest genetically characterized ASD cohort to date^71^. It is likely that hundreds of new *PTEN* mutation carriers will be identified in SPARK and would be available for recontact and detailed prospective study.

We demonstrate that comprehensively assaying the molecular phenotypes of thousands of mutations to a human protein can yield clinically relevant insights, even for proteins with pleiotropic effects. Future efforts that combine multiple functional modalities and rich clinical datasets may allow for the precision needed to fully realize personalized genomic medicine.

## METHODS

### *PTEN* saturation mutagenesis

Our mutagenesis approach was similar to the Mutagenesis by Integrated TilEs (MITE) approach^32^. We designed a series of “tiles” that were complementary to wild-type PTEN except for one codon (Supplementary Fig. 1a). At this single codon, each molecule bore a substitution to the yeast-optimized codon for each non-wild-type amino acid, the yeast-preferred stop codon, or an in-frame, single codon deletion. Additionally, each set of “tiles” contained unique DNA adapters on either end to allow PCR retrieval of individual tiles from the pool (using primers with prefix: PTEN_sliceprimer, Supplementary Table 1). These DNA “tiles” were synthesized as 130-mers as part of a 12,000-feature oligo pool by CustomArray (Bothell, WA prefix: PTENTile). For each “tile”, we designed inverse PCR primers that linearized the pYES2-PTEN wild-type sequence, excluding the portion encoded by the corresponding tile. Following amplification the “tile” PCR products were incorporated into the appropriate linear pYES2-PTEN by SLiCE mediated recombination^33^. SLiCE reactions were 10 μL and consisted of 100 ng of linearized vector with 15 ng of “tile” DNA, along with 1x SLiCE buffer and 1x SLiCE extract (SLiCE extract and buffer were prepared as described^72^). Reactions were incubated for 60 minutes at 37°C, then diluted 1:10 in water and 2.5 μL used to electroporate 50 μL of NEB 10-beta electrocompetent E. Coli. Transformation reactions were plated on LB agar plates with 100mg/mL Carbenecillin (GoldBio) and grown overnight at 37°C. Colonies were collected and plasmids isolated with the QIAprep Spin Miniprep Kit (Qiagen).

### Yeast selection experiments

Plasmid libraries were normalized and pooled into 4 mega-pools, each representing saturation mutagenesis for one quadrant (quadrants 1-3 = 100 codons, quadrant 4 = 103 codons). One μg of each mega-pool was transformed into the S. *cerevisiae* strain YPH-499 already containing YCpLG-p110μ-CAA☓ using the Li-Ac/SS carrier DNA/PEG method^73^ resulting in >50,000 colony forming units per reaction. Libraries were grown overnight in SC-glucose ‒leu ‒ura (synthetic complete medium lacking leucine and uracil, using glucose as carbon source), pelleted and frozen down in 15% glycerol in ‒80°C.

Selection experiments began with overnight outgrowth of frozen stocks in SC-raffinose ‒leu ‒ura (raffinose neither induces nor represses GAL1/10 promoter). Following outgrowth, 25 or 30 million cells (replicate A or B) were pelleted for each quadrant as the “input” sample and frozen at ‐20°C. Also, 25 or 30 million cells were seeded into each of (3) 50 mL SC-galactose ‒leu ‒ura, and (1) SC-raf ‒leu ‒ura. Cultures were incubated at 30°C with 185 rpm shaking. After 24 hours of growth, cell concentrations were measured with TC-20 Automated Cell Counter, and for each replicate 20 million cells were passaged into fresh medium. At 36 and 48 hours, the same was done, except that 20 million cells of each sample were also kept as a timepoint sample. Samples were spun down with 13,000 × g for 30 seconds, medium withdrawn, and frozen in ‐20°C.

### Library prep and sequencing

Plasmid DNA was isolated from pelleted cells with Zymoprep Yeast Plasmid Miniprep II kit (Zymo Research). Stage-one PCR to append partial Illumina adapters was performed on 5 ng DNA with primers pYES2-PTEN_Q[1-4][F/R]_S1 and using KAPA HiFi Hotstart Readymix (KHF) in 25 μL reactions (1× KHF, 0.5 uM each primer, 2.5 ng DNA, 1× SYBR Green). Reactions were monitored by qPCR with cycling conditions: [95°C 3 minutes (98°C 20 seconds, 55°C 30 seconds, 72°C 15 seconds, plate read, 72°C 8 seconds) × 28-36 cycles]. Reactions were removed during or immediately following exponential phase of amplification. Stage-two PCR was then done in 25 μL reaction volumes on 1 μL of uncleaned stage-one product using 0.5 μM of primers pYES2-PTEN_4Q_S2_F_[1-2] and pYES2-PTEN_4Q_S2_R_[1-44], 1× KHF, and 1× SYBR Green. Reactions were cycled with [95°C 3 minutes (98°C 20 seconds, 55°C 15 seconds, 72°C 15 seconds, plate read, 72°C 8 seconds) x 6 cycles]. Reaction products were checked on a 1.5% agarose gel, purified using NucleoSpin PCR Clean-up (Machery-Nagel), and concentrations were measured using Nanodrop. Samples were normalized and combined into a common pool that was sequenced across multiple runs using paired-end 300 base-pair reads on Illumina MiSeq platform (v3 Reagent kit).

### Sequencing data analysis

Paired end reads were merged with PEAR^74^ and common priming sequences were trimmed from the 5’ and 3’ ends using cutadapt^75^. For each quadrant, a purely wild-type sample was sequenced in order to identify the sequencing error profiles. Counts of error reads were normalized to wild-type counts, then this normalized amount of reads were removed from all experimental samples^7^. Sequence variants were identified and counted with custom python scripts. These raw variant counts files were analyzed with Enrich2 v1.2.0 (https://github.com/FowlerLab/Enrich2)^34^ to calculate scores and standard errors for each variant.

### Mutation collation

We collected ASD-associated variants from SFARI Gene^76^ (accessed 10/09/17) and the literature. We collected PHTS-associated mutations from the literature. A mutation was considered ASD/DD-associated if the report did not include symptoms of PHTS, and the mutation had not been reported in another individual with PHTS. If an individual had ASD/DD and PHTS features, or was observed in multiple individuals representing both presentations, we considered it PHTS. We considered any PTEN missense or nonsense (excluding frameshifts) mutation in the gnomAD database^1^ (accessed 11/19/17) to be benign, with the exception of two mutations that are considered pathogenic in ClinVar (K289E and R173H). We considered single-residue missense mutations from ClinVar (accessed 09/30/17) that were considered either pathogenic or likely pathogenic, were submitted with criteria, and had no conflicting reports to be pathogenic.

### Protein positional features and modeling

Conservation values were acquired from Consurf DB^77^ with default settings. Relative solvent exposure was calculated with GETAREA web tool^78^. For Figure 3, a position was considered exposed if its ratio of side-chain surface area to “random-coil” surface area exceeded 50, intermediate if the ratio was between 20 and 50, and buried if its ratio was less than 20. Secondary structure assignments were enumerated with STRIDE^79^. Pymol (href="https://pymol.org/2/) was used to generate structural representations with pdb entry 1D5R. Clustering was performed on the 326 positions with all 19 missense mutations measured (including high and low confidence). Clustering was performed with scipy.cluster.hierarchy.linkage, method= “ward”.

### Mutation effect predictors

We obtained Provean and SIFT predictions from Provean Protein (http://provean.jcvi.org/seqsubmit.php) with default settings. We considered “Deleterious” Provean predictions as pathogenic and “neutral” Provean predictions as benign. For SIFT, we considered “Damaging” predictions as pathogenic and “Tolerated” predictions as benign. We obtained Polyphen-2 predictions from Polyphen-2 batch query web server (http://genetics.bwh.harvard.edu/pph2/bgi.shtml). For Polyphen-2, “probably damaging” or “possibly damaging” predictions were considered pathogenic, whereas “benign” predictions were considered benign.

## ACKNOWLEDGMENTS

We thank R. Pulido for providing yeast expression constructs and YPH-499 yeast strain and Y. Zhang for providing *E. coli* strain used for generating the SLiCE reagent. We thank Y. Jia and the Oregon National Primate Research Center Molecular & Cell Biology Core for technical assistance. We thank A.C. Adey, E. Fombonne, D.M. Fowler, A. Rubin, J. Mester, J. Weile, J. Savage, U. Shinde, J. Zonana, G. Mandel, P. Stork, and K. Wright for helpful discussions. We thank Martha Atherton and the Atherton Foundation for their support of the NARSAD awards. This work was supported by the Brain and Behavior Research Foundation through the NARSAD-Atherton Foundation Young Investigator Award (59116 to B.J.O.) and internal funds (B.J.O.). T.L.M received support through the National Institutes of Health (NIH) Grant T32DK007680. T.L.M. is an ARCS scholar (Achievement Rewards for College Scientists Foundation, Inc., Oregon Chapter), B.J.O. is a Klingenstein-Simons Fellow (Esther A. & Joseph Klingenstein Fund, Simons Foundation) and Sloan Research Fellow in Neurosciences (Alfred P. Sloan Foundation, FG-2015-65608).

## AUTHOR CONTRIBUTIONS

B.J.O. and T.L.M. conceived and designed the study. T.L.M. and S.E. validated, modified, and optimized the mutagenesis and liquid culture conditions. T.L.M. created the mutation libraries. T.L.M. conducted the selection experiments and created computer scripts for data analysis. T.L.M. and B.J.O. analyzed the data and wrote the manuscript.

## COMPETING FINANCIAL INTERESTS

The authors declare no competing financial interests.

